# Prediction of fluorophore brightness in designed mini fluorescence activating proteins

**DOI:** 10.1101/2021.07.21.452510

**Authors:** Emma R. Hostetter, Jeffrey R. Keyes, Ivy Poon, Justin P. Nguyen, Jacob Nite, Molecular Modeling and Design Class, Carlos A. Jimenez Hoyos, Colin A. Smith

## Abstract

The de novo computational design of proteins with predefined three-dimensional structure is becoming much more routine due to advancements both in force fields and algorithms. However, creating designs with functions beyond folding is more challenging. In that regard, the recent design of small beta barrel proteins that activate the fluorescence of an exogenous small molecule chromophore (DFHBI) is noteworthy. These proteins, termed mini Fluorescence Activating Proteins (mFAPs), have been shown increase the brightness of the chromophore more than 100-fold upon binding to the designed ligand pocket. The design process created a large library of variants with different brightness levels but gave no rational explanation for why one variant was brighter than another. Here we use quantum mechanics and molecular dynamics simulations to investigate how molecular flexibility in the ground and excited states influences brightness. We show that the ability of the protein to resist dihedral angle rotation of the chromophore is critical for predicting brightness. Our simulations suggest that the mFAP/DFHBI complex has a rough energy landscape, requiring extensive ground-state sampling to achieve converged predictions of excited-state kinetics. While computationally demanding, this roughness suggests that mFAP protein function can be enhanced by reshaping the energy landscape towards states that better resist DFHBI bond rotation.

## Introduction

The creation of *de novo* designed proteins is a burgeoning area of structural biology that opens a vast range of new protein geometries. However, giving those designs useful functions can sometimes be considerably more difficult than making them correctly fold. In that regard, the recent *de novo* design of mini Fluorescence Activating Proteins (mFAPs)^1^ that bind to and activate the fluorescence of a small molecule chromophore is particularly noteworthy. The design process involved creating β-barrels from scratch then modifying those structures to harbor a chromophore inside the barrel, producing proteins somewhat analogous to green fluorescence protein (GFP)^2^, albeit half the size. The designs used the 3,5-difluoro-4-hydroxybenzylidene imidazolinone (DFHBI) chromophore, an exogenously synthesized mimic of the GFP chromophore previously created for the SPINACH fluorescent RNA aptamer^3^. It is composed of a fluorine substituted phenolate ring and an imidazolinone ring joined by a methine bridge with conjugated bonds. While DFHBI shows negligible fluorescence in solution, its fluorescence increases dramatically when sandwiched between RNA bases in SPINACH^4^ or enclosed in the binding pocket of the mFAP designs. After creation of the initial mFAP/chromophore complex, other variants were designed and tested, improving on both the chromophore binding affinity and quantum yield.

One of the best designs, mFAP2^1^, was subsequently refined^5^ to show further increases brightness through an iterative series of combinatorial library screens. While efforts were made to improve packing and chromophore affinity both by manual and computational selection of proposed mutations, the primary means of improving function came through experimental screening of combinatorial libraries instead of through a mechanistic model of the fluorescence process. In addition to improving brightness, the authors also designed a variant, mFAP_pH, that showed similar binding affinities to the negatively charged (phenolate) and neutral (phenolic) forms of DFHBI, functioning as a pH sensor due to the respective red- and blue-shifted excitation wavelengths of the two DFHBI charge forms. Furthermore, the authors created a Ca^2+^-sensor by fusing calcium binding domains to the mFAP, which then showed increased DFBHI binding affinity in the presence of Ca^2+^, enabling switching via a chromophore bind and release mechanism. This cooperative binding mechanism was recapitulated in molecular dynamics simulations showing that Ca^2+^ binding shifted the mFAP towards conformations similar to those observed with DFHBI bound.

While addition of the pH and Ca^2+^ sensing modalities could be computationally evaluated using hypothesized mechanisms of action, optimization of chromophore brightness relied almost entirely on library design and screening. This is notable because the highest quantum yield for any mFAP design was 0.237^1^, quite a bit lower than GFP (0.79)^6^, EGFP (0.60)^6^, or SPINACH (0.72)^3^. Efforts to make further improvements in mFAP fluorescence should be greatly assisted by a mechanistic understanding of the determinants of fluorescence, and the ability to computationally simulate that mechanism.

The photophysics of DFHBI bound to SPINACH has been an active area of research^3,7–9^. When exposed to continuous light at the same illumination level, RNA-bound DFHBI exhibited less photobleaching than EGFP or fluorescein^3^. Time-resolved experiments showed a single exponential fluorescence decay with a lifetime of 4.0 ns^7^, slightly longer than the GFP lifetime of 2.7 ns. Fluorescence recovery after photobleaching from high intensity light was found to primarily occur through exchange of dark state chromophores with other free DFHBI molecules^7^. Several simulation studies have investigated the role of SPINACH in promoting fluorescence. Quantum mechanics/molecular mechanics (QM/MM) point calculations from energy minimized structures have suggested that the presence of SPINACH increases the barrier to rotation of the imidazole group primarily through interaction with the MM environment and not through RNA-induced perturbation to the QM energies^8^. A subsequent study highlighted the role of the electrostatic environment in shifting chromophore vertical excitation energies^9^.

The GFP chromophore upon which DFHBI was based has been very well studied. Like DFHBI, the GFP chromophore has a phenolate group attached to an imidazole group by a methine bridge. However, the imidazole group is covalently attached to the protein backbone because the chromophore is autocatalytically formed from several residues of the GFP protein^10,11^. Chromophore rotation has been of considerable interest, allowing both non-radiative relaxation back to the ground state as well as reversible photoswitching between dark/light states of GFP and GFP-like proteins^12–15^. Several contrasting rotation pathways have been proposed for photoisomerization of chromophores in general, including the one bond flip (OBF)^16^, in which rotation occurs primarily via one bond, and the hula-twist (HT)^17^, in which rotation occurs via adjacent bonds in a concerted fashion. In GFP, the charge state and electronic distribution within the chromophore has been shown to have a significant impact on the energetics of each pathway^18^.

Several studies have used molecular dynamics simulations to simulate the rotation of the methine bridge dihedral angles in the GFP chromophore. The first to do so modeled the dihedral angles of the methine bridge as being freely rotatable in the excited state^13,19^, with a flat dihedral angle potential. The amount of dihedral freedom was found to correlate with the quantum yield across four GFP-like systems. Park and Rhee^20^ used a significantly enhanced model of chromophore energetics to simulate a ground state (S_0_) equilibrium ensemble, followed by excited state (S_1_) simulations, in which the probability of making a non-radiative transition back to the ground state was determined with Landau-Zener surface hopping^21^. The energy of the chromophore was determined using an interpolated mechanics/molecular mechanics (IM/MM) approach, where energies for the chromophore were determined by interpolating quantum mechanical energies from 1500 chromophore conformations. They found that the electrostatic potential, especially around the phenolate oxygen, was particularly important for determining the kinetics of chromophore rotation and nonradiative decay.

Here we use a combination of quantum mechanics and molecular dynamics simulations to model how several mFAP designs keep the DFHBI chromophore planar after excitation. We find that relatively simple quantum mechanics calculations can produce potential energy surfaces that enable molecular dynamics simulations to qualitatively distinguish between brighter and dimmer mFAP variants. We focus mainly on pairs of mFAP designs differing at a single amino acid position, finding that differences in brightness can be more accurately predicted for single mutations than multiple mutations. We also find that the mFAP conformation at the time of excitation has a large influence on its ability to restrict the chromophore to a planar conformation.

## Results and Discussion

### Quantum Mechanical Calculation of Potential Energies

The ability of a host macromolecule to hold DFHBI and other GFP-like chromophores in a planar conformation has long been understood to be a major contributor to fluorescence, so we first sought to quantify how intrinsically resistant DFHBI was to dihedral rotation. To do so, we used gas-phase density functional theory (DFT) quantum mechanical (QM) calculations to determine the change in potential energy when either the imidazole or phenyl dihedral angles (Figure 1A) were rotated away from a planar cis conformation. We first analyzed the ground state and found that the planar conformation (having each dihedral defined as 0°) was very rigid, with large energy barriers at 90° for both the imidazole (sometimes referred to as τ, 188 kcal mol^-1^) and phenyl (φ, 129 kcal mol^-1^) dihedrals (Figure 1B) The imidazole potential energy surface was similar to that determined a previous study using MS-CASPT2 calculations^8^, which reported a barrier of about 170 kJ/mol. Notably, the dihedral potential energies determined from QM were 6-7 times larger than those generated from Generalized Amber Force Field (GAFF)^22^ parameterization, suggesting that conjugation between the rings was not entirely accounted for by the GAFF parametrization tools.

**Figure 1.**
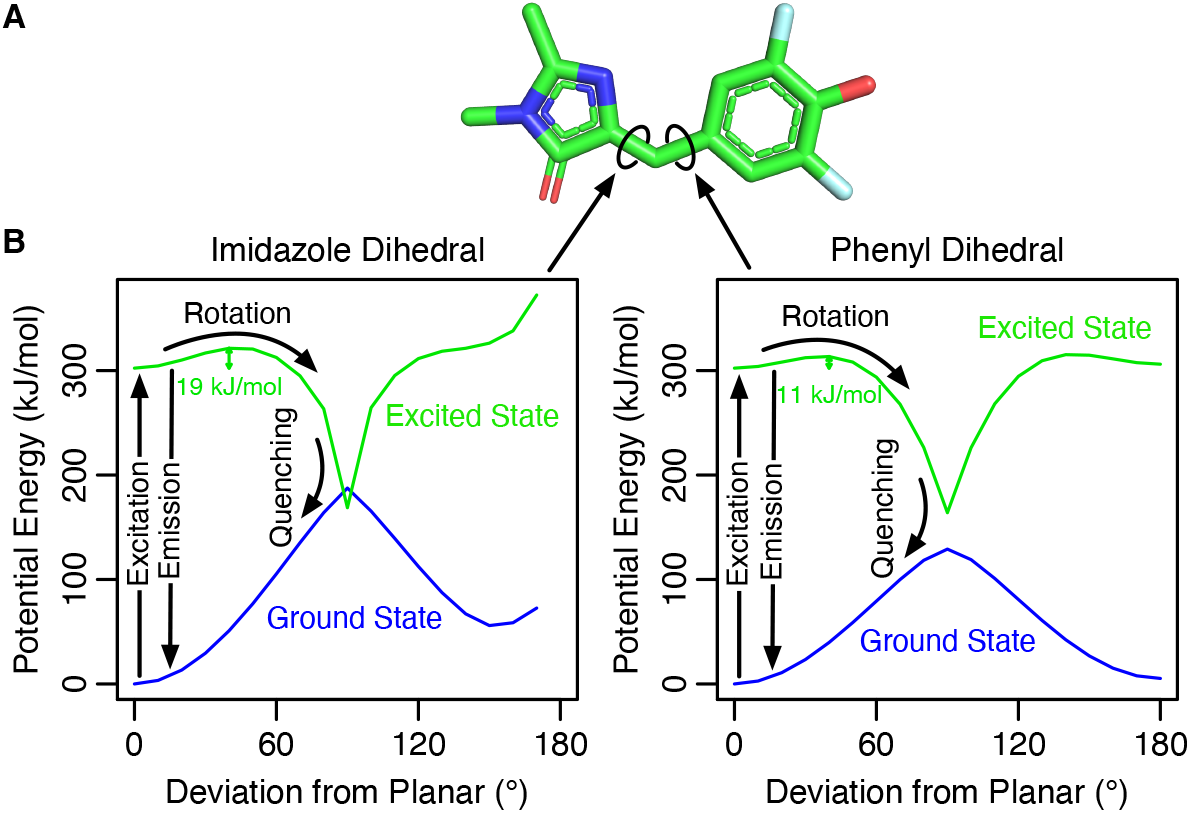
Model for DFHBI quenching upon dihedral rotation. **A)** DFHBI has two rotatable bonds associated the imidazole and phenyl groups. **B)** Gas-phase quantum mechanical calculations of the ground state (blue) show that both dihedral angles are energetically predisposed to stay planar, with high (>100 kJ/mol) barriers at 90°. However, in the excited state (green), the planar conformation is only metastable. The proposed model of quenching involves competition between photon emission and rotation followed by a non-radiative transition back to the ground state.

Due to structural symmetry, the phenyl torsional profile was symmetric, whereas the imidazole was not. The lowest energy alternate conformation of the imidazole (close to trans at 140°) was 56 kJ mol^-1^ higher than the planar state due to a steric clash between the carbonyl oxygen on the imidazole and a hydrogen on the phenyl ring. In GFP-like proteins, 180° rotation of the imidazole dihedral is known to cause photoswitching^14^. It has been proposed that cis/trans isomerization of SPINACH-bound DFHBI about the imidazole dihedral is also responsible for the dark state observed after excitation^7^. Prior to excitation, DFHBI molecules capable of fluorescing should therefore exist in a single dominant conformational state (shown in Figure 1A) at room temperature, with only small deviations from planarity.

With DFHBI being largely inflexible in the ground state, we used time-dependent density functional theory (TD-DFT) QM to calculate potential energy surfaces for the first singlet electronic excited state. While it has been noted that TD-DFT can overestimate the vertical excitation energies of molecules like DFHBI^8^, it was also found that the method used to optimize the geometry had little effect on the excitation energy, indicating that TD-DFT can provide reasonable excited state geometries. Here, we were primarily interested in the energy barrier to rotation rather than the excitation energy, so TD-DFT should provide reasonably accurate geometries without having to resort to more expensive methods. We found that while the planar conformation in the ground state was both kinetically and thermodynamically stable, the planar conformation was only metastable (i.e. a local minimum) in the excited state. There was an estimated 19 kJ mol^-1^ (imidazole dihedral) or 11 kJ mol^-1^ (phenyl dihedral) energy barrier between the planar conformation and the minimum potential energy at 90°. (Figure 1B) This higher energy barrier for the imidazole dihedral also agreed with a previous study that used MS-CASPT2 QM simulations^8^. For both dihedrals, the excited state energy either intersected with or approached the ground state energy at 90°. However, the level of QM theory (DFT and TD-DFT) used here is insufficient to fully capture energetics in that region of phase space.

These energy surfaces are consistent with the hypothesis that once excited, there are two dominant pathways for return to the ground state: either emission of a photon, or dihedral rotation over either energy barrier followed by non-radiative relaxation, resulting in quenching of fluorescence. (Figure 1B) In this model, even if there is not a true conical intersection, the difference in energy between the ground and excited states at an angle close to 90° will likely be too small to result in emission of high energy photons. This indicates that the primary function of the mFAP proteins is to increase the kinetic barrier between the metastable planar state and the non-emissive rotated state, with the different amino acid sequences affecting the height of the barrier and ultimately the fluorescence. This is similar to the proposed mechanism for Spinach-DFHBI, where the majority of the energy barrier to dihedral angle rotation comes from non-quantum interactions between the chromophore and DNA bases^8^.

### Simulation of DFHBI Photon-Assisted Rotation and Quantum Yield

As an initial test of this hypothesis, we ran simulations of free DFHBI in water to determine whether the observed rotation kinetics would quantitatively agree with experimental measurements of the quantum yield. While hybrid quantum mechanics/molecular mechanics (QM/MM) simulations would ordinarily be suited for this purpose, their computational cost would have been prohibitive for accumulating enough statistics to obtain converged estimates of the time it takes to cross the energy barriers, especially for the long rotation times expected in the context of a protein. Instead, we chose to use the gas-phase QM potential energies to reparametrize GAFF molecular mechanics (MM) parameters for the two dihedral angles. For both the ground and excited states, we found that potential energies for either transition pathway (0-80°) could be very well approximated using a series of cosine functions. (Figure S1) This method of parameterization makes the approximation that the energies of the dihedrals are independent of one another and can be treated in an additive manner. To determine whether that has an impact on the most sampled regions of the potential energy surface, we calculated a full 2D potential energy surface of both dihedrals sampled simultaneously. We found little deviation in the metastable region between the full non-additive model and an additive model similar to the one used in the molecular mechanics-based simulations. (Figure S2)

To determine the distribution of excited state rotation times for DFHBI in water, we first generated a Boltzmann ensemble of ground state conformations using 100 ns simulations with the ground state potential. (Figure 3A) Snapshots were saved every 50 ps and used to start new excited state simulations in which the nuclear coordinates and velocities were preserved, but the dihedral angle potential was instantaneously switched to the excited state parameters. All other potential parameters were the same as in the ground state. From there, both the imidazole and phenyl dihedral angles were monitored every 50 fs and DFHBI was considered to have rotated/quenched when either angle exceeded 60°. Using that data, we calculated a decay curve showing the fraction of DFHBI simulations remaining planar as a function of time since excitation. (Figure 2A) A fit to that curve indicates that the planar state of DFHBI shows a purely single-exponential decay with a rate constant *k*_planar_ = 0.131 ps^-1^ and average lifetime τ_planar_ = 7.6 ps when simulated in this environment.

**Figure 2.**
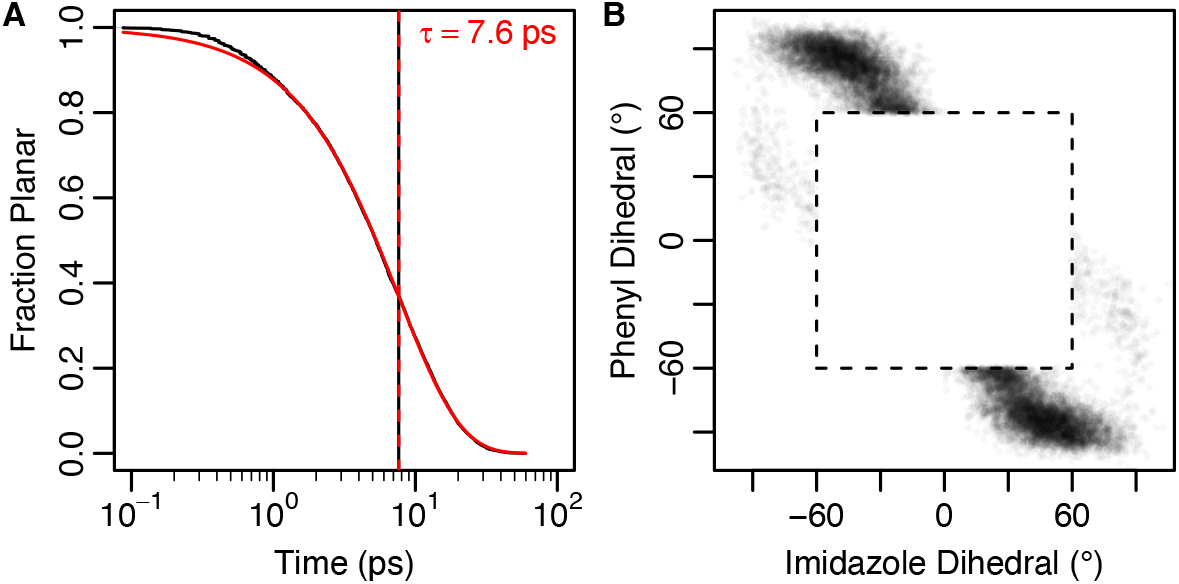
DFHBI in water shows monoexponential escape from planar geometry. **A)** When simulated in water with 150 mM NaCl, the planar state shows a monoexponential decay (red) with average lifetime of 7.6 ps. A subtle lag is visible in the first picosecond. Escape from the planar state was defined as the time at which either the imidazole or phenyl dihedrals exceeded 60°. The decay curve (black) and standard error (shaded gray) is calculated from four independent 100 ns ground state simulations and yields the same average lifetime (vertical black line). **B)** The first recorded set of dihedral angles for each trajectory after leaving the planar state (dashed lines). Simulation snapshots were recorded every 50 fs, so there is up to a 50 fs time lag between leaving the planar state and recording of the dihedral angles. Most trajectories escaped the planar region via the phenyl dihedral transition state.

**Figure 3.**
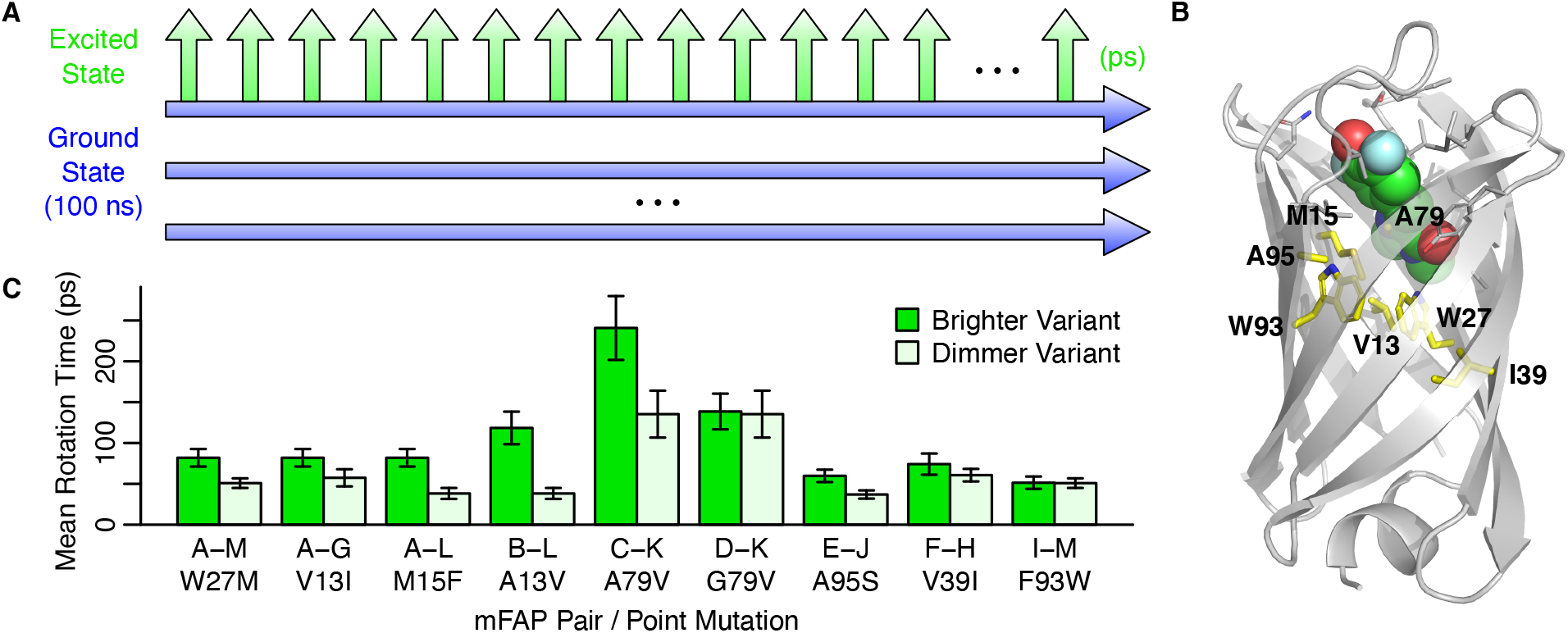
Molecular dynamics simulations predict effects of point mutation on fluorescence. **A)** 20 100-nanosecond ground state simulations (blue) were started from slightly different ligand poses, with excited state simulations (green) branched off every 50 picoseconds. The rotation time (typically picoseconds) was recorded when one of the two DFHBI dihedral angles exceeded 60 degrees. **B)** Mutations at seven different amino acid positions were compared, shown here in yellow with the sequence from mFAP-A. **C)** In all cases, the brighter variant showed a mean rotation time that was greater than or equal to the dimmer variant. For each of the ground state simulations, the average rotation time over the 2000 excited state simulations was calculated. Error bars give the standard error of the mean starting from 20 different ground state simulations.

For quantitative comparison with experimentally measured extinction coefficients, we used the observed fluorescence lifetime of Spinach-DFHBI (τ_fluor,obs_), which was previously reported to be 4.0 ns^7^. If that fluorescence lifetime is normalized to account for the observed DFHBI-Spinach quantum yield (QY) of 0.72^3^, then the underlying fluorescence lifetime assuming no quenching occurs is τ_fluor_ = τ_fluor,obs_/QY = 5.56 ns. If that is assumed to be the fluorescence lifetime of DFHBI given that it is in the planar state (τ_fluor|planar_), then the quantum yield predicted by the simulations would be τ_planar_/(τ_fluor|planar_ + τ_planar_) = 0.0014, which is only two-fold greater than the measured value of 0.0007^23^. This suggests classical molecular dynamics simulations using GAFF parameters augmented with dihedral angles reparameterized from DFT/TD-DFT QM energies may be sufficient to approximate the atomic kinetics of the DFHBI ground and excited states.

As expected from the lower energy barrier, most trajectories escaped the metastable planar state through rotation of the phenyl dihedral. (Figure 2B) However, the motion was not purely a one bond flip, as very few trajectories exited the planar state with the other dihedral angle still close to 0°. Instead, the simulations showed a hybrid between the one bond flip and hula-twist mechanisms, with an average deviation from planar of 20-25° for one angle at the moment the other angle exceeded 60° from planar.

### Prediction of mFAP Variant Brightness

To test the hypothesis that mFAP variant fluorescence could be explained by the ability of each sequence to resist DFHBI dihedral rotation, we identified pairs of variants having a single amino acid difference and greater than two-fold change in normalized fluorescence (see Methods and Figure S3). Each of the 13 studied variants was given a letter corresponding to the magnitude of the fluorescence, with mFAP-A being the highest and mFAP-M being the lowest. Across all pairs, there were seven different residue positions with single point mutations, most being part of the hydrophobic core packing against DFHBI. (Figure 3B) Several of the mutated positions did not make direct contact with DFHBI, including residues 39, 93, and 98.

For each variant, we ran the same simulation protocol that was used for free DFHBI (Figure 3A), starting 20 independent ground-state simulations from slightly different ligand poses and side chain conformations. For each ground-state simulation, a set of 2,000 individual excited-state trajectories was run. We calculated the average time it took the chromophore to rotate out of the planar state for each set. The overall rotation statistics for a given variant were determined by taking the mean and standard error over the 20 per-set average rotation times. According to the hypothesis, the brighter variant out of each pair should have a longer average rotation time, which was largely borne out by the data. When the point-mutant pairs were compared, every one of the more fluorescent variants had an equal or greater mean rotation time than the corresponding dimmer variant. (Figure 3C) Three of the pairs showed overlapping error bars, with two out of the three having low fluorescence for both proteins in the pair: mFAP-F/H and mFAP-I/M. The other pair was mFAP-D/K, which is discussed in more detail below.

For all 13 variants, we then compared the experimental normalized fluorescence values to the mean rotation times and found a positive correlation (R = 0.56, Figure 4), indicating that a sizable fraction of the variation in fluorescence can be explained using dihedral rotation kinetics derived from the MD simulations. As expected from the comparison of pairs in Figure 3C, lines connecting variants differing by a single amino acid all showed a positive slope.

**Figure 4.**
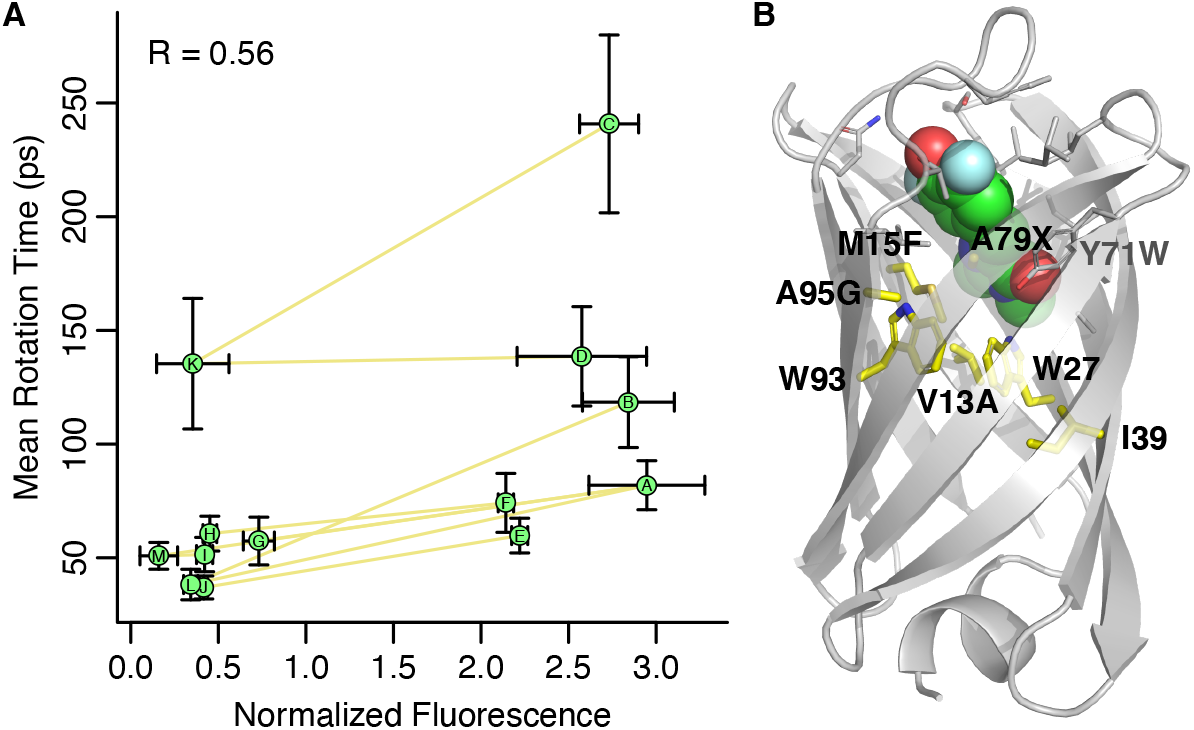
Chromophore rotation time shows correlation with fluorescence. **A)** Sequence variants differing by a single amino acid position are connected by yellow lines. mFAP-K is a possible outlier because its binding affinity (~60 μM) was at least tenfold weaker than all other variants. **B)** A cluster of related sequences differing at position 79 (mFAP-C/D/K) show overall longer rotation times, likely due to four additional background mutations with respect to mFAP-A: V13A, M15F, Y71W, and A95G. (Figure S5) mFAP-B shares two of those mutations: V13A and M15F.

One of the variants, mFAP-K, showed the third longest mean rotation time but low normalized fluorescence. That discrepancy may arise because the DFHBI binding affinity for mFAP-K (~60 μM) was an order of magnitude weaker than any other variant. (Table S2) A saturating ligand or protein concentration was therefore less likely, especially at the DFHBI concentrations used in the fluorescence assays (0.1 and 10 μM). That probably led to a lower concentration of the actively fluorescent mFAP-K/DFHBI complex, thus reducing the observed fluorescence. If this was a real effect, mFAP-K fluorescence should be the most sensitive to the DFHBI concentration out of the variants considered. Indeed, the mFAP-K fluorescence at 0.1 μM DFHBI was far lower relative to 10 μM DFHBI than any other variant, supporting the role of concentration/binding affinity affecting observed fluorescence. (Figure S4) It is reasonable that incomplete binding may also attenuate fluorescence at 10 μM DFHBI. Thus, mFAP-K brightness may be significantly higher than the normalized fluorescence suggests. When mFAP-K is removed, the correlation coefficient increases to 0.68.

Another source of error may come from background mutations that together contribute to artificially longer or shorter simulated DFHBI dihedral rotation times. For instance, a cluster of three variants, mFAP-C/D/K, showed the longest rotation times. In addition to having different amino acids at position 79 (alanine/glycine/valine, respectively), those four variants differ from mFAP-A by four additional background mutations. (Figure 4B and Figure S5) Small errors introduced by each of those mutations may accumulate and result in rotation times that were either too long for mFAP-C/D/K, or too short for the other mFAP variants. The gradual accumulation of bias is supported by mFAP-B, which shares two of the four background mutations and has a rotation time situated between mFAP-C/D/K and the other variants.

While the mFAP-C/D/K cluster may have shown higher than expected rotation times, the effects of multiple mutations in another cluster, mFAP-E/F/J/H, were predicted correctly. That cluster differed from mFAP-A at least five amino acid residues: 67, 100-102, and 104. None of those side chains made direct contact with DFHBI in mFAP-A, yet they had the overall effect of slightly reducing the experimentally measured fluorescence of mFAP-E/F relative to mFAP-A. This moderate reduction was mirrored in the rotation times of mFAP-E/F. Put together, these results suggest that while the effects of multiple mutations can be captured in some cases, the most predictive application of the simulations is stepwise, single amino acid changes.

### Compensatory and Allosterically Acting Mutations

While the effects of multiple mutations were sometimes less accurate than point mutations, the simulations were notable in their ability to capture compensatory, non-additive effects. For instance, the variants studied here contained a complete “evolutionary” pathway going from mFAP-A to mFAP-B, one point mutation at a time. (Figure S5) While those two variants had similar fluorescence, the intermediate mFAP-L had much lower fluorescence. The mean rotation times predicted by the simulations mirrored those results, with mFAP-A/B having significantly longer rotation times than mFAP-L. Going from mFAP-A to mFAP-L involves substituting methionine 15 for a phenylalanine, which is slightly larger and able to form stacking interactions with DFHBI, but decreases its fluorescence. Fluorescence is restored, both experimentally and computationally, by mutating the adjacent valine 13 to smaller alanine. Interestingly, while the MD simulations were able to predict lower fluorescence for mFAP-L, its measured binding affinity (1.9 μM) was very similar to mFAP-A (1.8 μM)^5^. Even if the binding affinity could be correctly predicted, it would not have been a good indicator of function.

In addition to correct modeling of those compensatory mutations, the simulations were able to capture the kinetic effects of mutations distant from the DFHBI binding site. For instance, the A95S mutation is 7-8 Å away from DFHBI (Figure 3B), but still results in a large reduction in fluorescence. The simulations likewise show a sizable reduction in the mean rotation time. A95 is in the beta strand immediately before loop 7, which extends over the top of DFHBI and may be important for maintaining that loop in a conformation that impedes DFBHI dihedral rotation.

### Effects of Ground State Flexibility and Excited State Energy Barrier

Preliminary simulations for this study were generated by a mixture of undergraduate and graduate students in the Molecular Modeling and Design course taught by the last author. The students generated starting structures with Rosetta, ran five replicate 100 ns ground state simulations, and 100 excited state simulations per ground state simulation. To facilitate rapid parameterization of the dihedral angles for the course, the geometries for quantum mechanical energy evaluation were generated by relaxing the structure in the ground state, followed by rigid rotation of the dihedral angles. The resulting energies and DFHBI parameterization were sufficient to produce the results described thus far.

To quantify how relaxation of the nuclear coordinates during the QM calculations affected the ground state energies, we ran additional DFT calculations, constraining the imidazole and phenyl dihedrals but allowing all other degrees of freedom to relax. The resulting 1D potential energy surfaces are shown as dashed lines in Figure 5A and Figure S6A. Relaxing the geometries resulted in a dihedral angle potential that was slightly more flexible, with the first cosine force constant parametrized for both dihedral angles being 76-77% of the value from rigid rotation parameterization.

**Figure 5.**
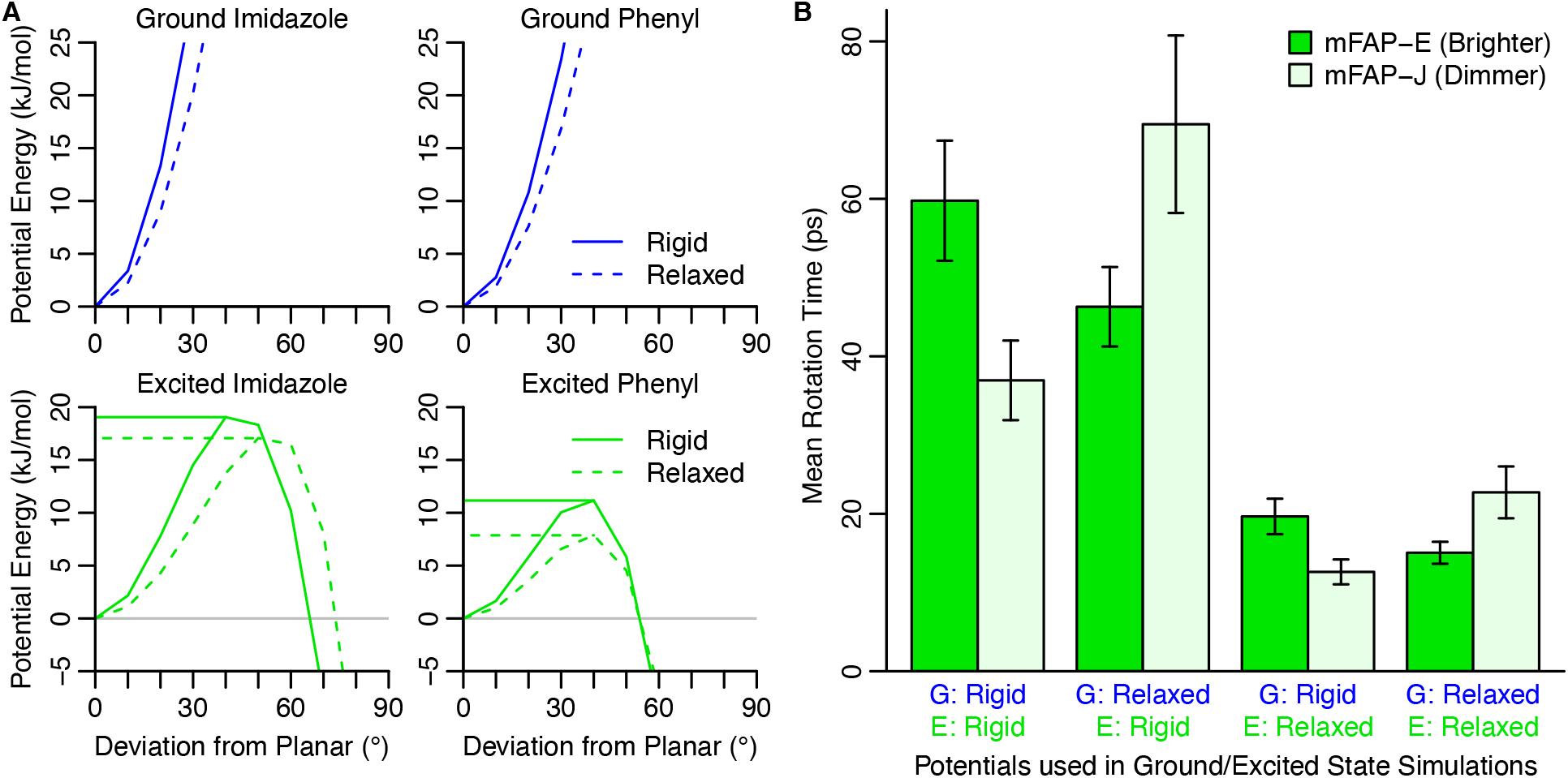
DFHBI ground-state potential shows sequence-specific effects on rotation times. **A)** Gas-phase quantum mechanics energies calculated using rigid dihedral rotation of the optimized planar geometry (solid lines) and relaxed geometry for each set of constrained angles (dashed lines). Excited state energies come from the same geometries as the ground state. Solid lines show the same data as Figure 1. **B)** Switching from the rigid (left set of bars) to relaxed (right set) dihedral potentials inverts predictions for the mFAP-E/J pair, unlike most other pairs which show the same predicted ranking (Figure S7). Mixing of the potentials (middle two sets) indicates that the inversion is primarily due to the ground state simulations. For mFAP-E/J, the different excited state potentials mainly result in scaling the overall rotation times.

We used those relaxed geometries to calculate vertical excitation potential energy surfaces for the first excited state. The energy barriers for imidazole and phenyl group rotation were reduced by 2-3 kJ/mol each over values from rigid geometric rotation. (Figure 5A and Figure S6A) While the phenyl energy barrier remained at approximately 40°, the location of the imidazole energy barrier increased to 50°. When the coordinates were relaxed in the excited state, the excited state energy barriers were further reduced by 6-11 kJ/mol from the ground relaxed coordinates, bringing them down to 5.7 and 2.1 kJ/mol for the imidazole and phenyl dihedrals, respectively. (Figure S6A) In addition, the location of the barrier was shifted from 50-60° to 20-30°, making jumps over the barrier much more probable.

To determine how these lower energy barriers affected the MD rotation times, we repeated the ground state simulations with the reparametrized potential. We used those same ground state simulations to launch two types of excited state simulations using either the ground-relaxed or excited-relaxed potential energy surfaces. The resulting rotation times for the original rigid and the two new relaxed excited state potentials are shown in Figure S7. As expected, the overall rotation times go down with decreasing excited state energy barrier, with overall average rotation times of 88 ps, 26 ps, and 0.58 ps, respectively. The brighter variant had the longer rotation times for most pairs in all three methods, suggesting that the discriminatory power of the MD simulations is not highly dependent on the barrier height in the excited state. The higher energy barrier from the original rigid potential did give a slightly higher correlation coefficient (R = 0.56 vs. 0.44 and 0.48) and correctly ranked the mFAP-E/J pair, while the relaxed potentials did not. (Figure 5B and Figure S7)

The mFAP-E/J pair showed one of the largest experimental differences in brightness (Figure S4 and Table S2), making it most likely to be an incorrect prediction for the relaxed potentials. To establish whether reversal of the mFAP-E/J rotation time ranking was because of the ground state or excited state simulations, we ran simulations with mixed combinations of the ground state and excited state potentials (i.e. rigid potential in the ground state and relaxed potential in the excited state). (Figure 5B) Intriguingly, the ranking of the excited state rotation times depended exclusively on which ground-state simulations were used, while the excited state potential served only to scale the overall rotation times. This may occur because for some protein sequences proximity to the energy barrier prior to excitation has a stronger impact on the observed rotation times. A related explanation, informed by observations in the next section, is that having a more flexible DFHBI molecule in the ground state may push the protein towards conformations that are less able to restrict DFHBI to a planar geometry in the excited state. In other words, a DFHBI molecule with more entropy in the ground state may carve out a reshaped binding pocket that yields additional conformational freedom in the excited state.

In an attempt to account for the non-additivity of the dihedral angle energies, we also calculated full 2D potential energy surfaces using the relaxed geometries. (Figure S8). We used those energies to calculate a grid-based correction term, which we applied in Gromacs using the CMAP functionality. The initial results from that work showed unexpectedly longer overall rotation times using the excited-potential energy surface. (Figure S9) This could result from differences in our grid spacing (10° vs. 15° for typical CHARMM CMAP potentials) or other implementation details that may have caused simulations artifacts. For instance, while a single CMAP correction is usually applied to pairs of adjacent bonds in the protein backbone, we applied two overlapping but identical CMAP corrections to account for all possible ways of forming the imidazole and phenyl dihedrals.

While our original QM calculations assumed rigid geometric rotations around the methine bridge dihedrals, they produced results that were somewhat more discriminatory between bright and dim mFAP variants than those using relaxed geometries. In addition to brightness, absolute quantum yields have been determined for DFHBI bound to mFAP-A (0.093) and mFAP-B (0.060)^5^. Whereas our simulations in water gave quantum yields two-fold greater than the experimentally measured value, the predicted quantum yields using the rigid QM energies for mFAP-A (0.014) and mFAP-B (0.020) were 3-7 fold lower than the experimental values. (Figure S10). Quantum yields from the ground relaxed QM energies for those two proteins were 10-23 fold lower than the experimental values.

Put together, these results suggest that either the true DFHBI dihedral angle potentials should be closer to the rigid ones, the Amber force field insufficiently models the proteins’ ability to restrain DFHBI rotation, or that something else is missing in the mFAP/DFHBI interactions. Modeling how the DFHBI electrostatic potential changes between the ground and excited states is one effect we plan to explore in future work. Indeed, it has been suggested that electrostatics play a dominant role in GFP chromophore dynamics^20,24^. Another effect we will investigate is the influence of the mostly apolar protein environment on the DFHBI potential energies, which contrasts with the more solvated GFP chromophore.

### Ground State Dynamics and Rotation Kinetics

While the DFHBI rotation kinetics in water were relatively simple, our simulations indicate that rotation becomes considerably more complex when hosted by an mFAP. In contrast to the aqueous simulations which showed monoexponential decay kinetics, binding to the mFAP variants showed biexponential or triexponential kinetics, indicating the presence of multiple rate constants. (Figure S10) Furthermore, to obtain sufficiently converged estimates of the mean rotation time, the results from 20 different ground-state simulations (each with a set of 2,000 excited-state simulations) were pooled together for each variant. This number was required because of considerable variability in the average rotation time between sets from the same variant. (Figure 6A) Depending on the variant, there was 7-49 fold difference between the shortest and longest average rotation times (i.e. the highest and lowest light green points in Figure 6A), each of which were derived from a different ground state simulation. The high degree of variability indicates that 100 ns is completely insufficient for an individual simulation to fully sample the Boltzmann ensemble of states that collectively determine the average rotation time. The individual ground state simulations are therefore likely sampling different parts of a rough mFAP/DFHBI energy landscape.

**Figure 6.**
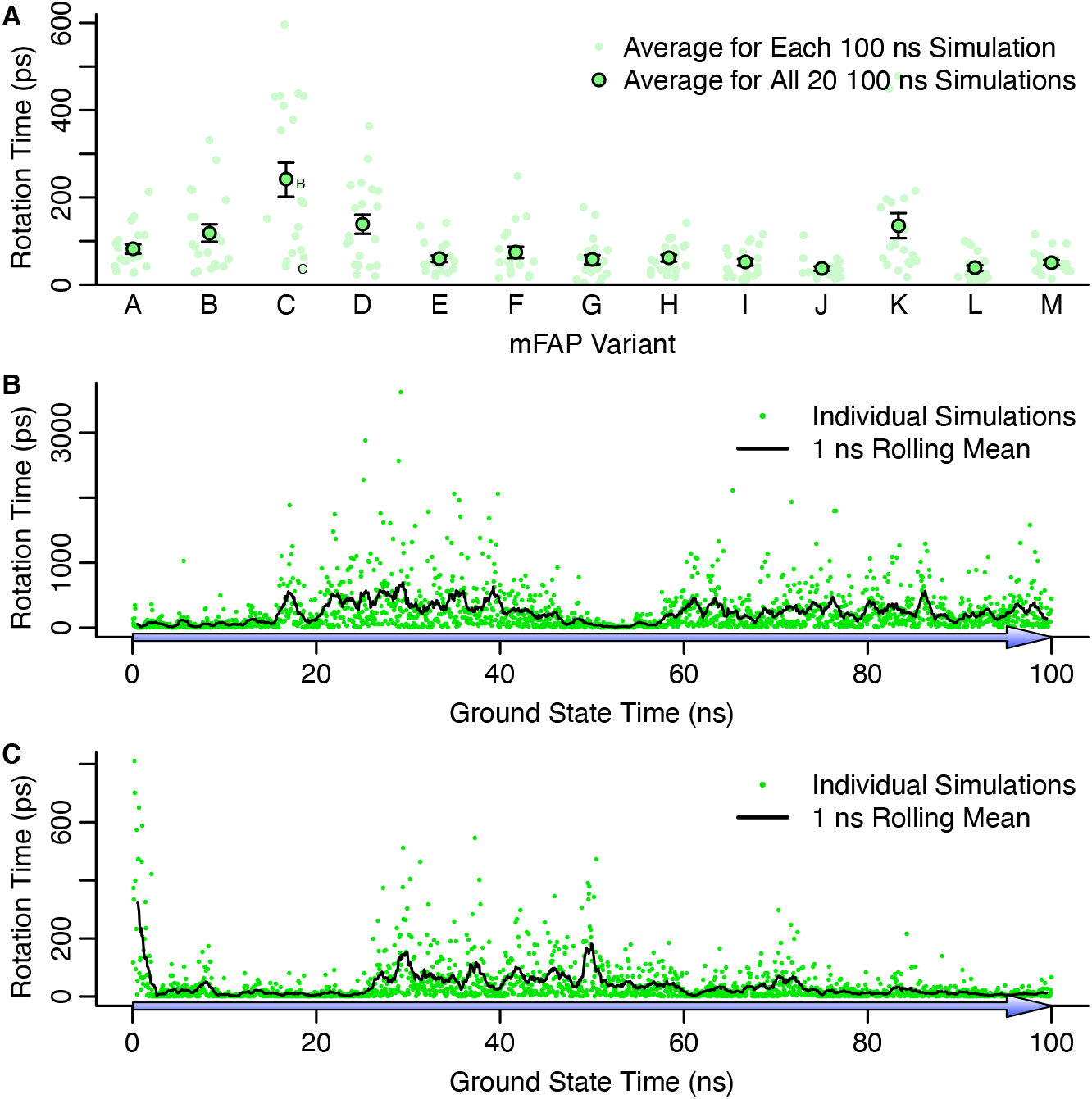
Large rotation time variability suggests a rough ground-state energy landscape. **A)** Light green points give averages over the 2000 excited state simulations branched from a single ground state simulation. Depending on the variant, there was a 7- to 49-fold difference in the longest average rotation time and the shortest average rotation time. The dark outlined points give the average over the 20 ground state simulations. These values/error bars are the same as shown in Figure 3C. **B)** and **C)** Within individual ground state simulations, there was also considerable variability in the rotation times. Although the raw data (green) was quite noisy, the 1 nanosecond rolling mean (black) shows long lived (tens of nanoseconds) states with longer rotation times, suggesting a change in structure. Simulations corresponding to **B** and **C** are labeled accordingly in **A**.

This high degree of variability also exists within individual ground-state simulations. Figure 6B/C shows all the excited-state rotation times for two of the mFAP-C ground state trajectories, one with the shortest average rotation time, and one that was very close to the overall mFAP-C rotation time. In these representative ground state trajectories, there were periods on the order of tens of nanoseconds long, in which excited state simulations showed much longer rotation times. The alternating trends can be more clearly seen in the 1 ns rolling mean of the observed rotation times. The observed heterogeneity and implied roughness of the energy landscape underscores the biophysical and computational complexity of this system. This suggests a model where the mFAP/DFHBI complex moves between different macrostates, each with distinct DFHBI rotation kinetics and thus different instantaneous brightness. While this makes prediction of the mFAP properties more complicated, it also suggests a rational approach for improving fluorescence by finding mutations that reshape the free energy landscape towards macrostates that show longer rotation times. We are currently in the process of trying to identify structural features that enhance fluorescence and design mutants that stabilize them.

## Conclusions

The prediction of protein function and activity is a longstanding goal of structural biology. Some functions can be predicted from thermodynamics, namely folding, ligand binding, and even allosteric regulation. However, other functions depend on the kinetics of non-equilibrium states, which can be more difficult to elucidate. There remain many challenges, especially describing systems in electronically excited states. This study highlights the critical role that conformational dynamics plays in determining the fluorescence of a chromophore bound to host proteins. In particular, while electrons are critical for absorbing and emitting photons, the kinetics associated with the nuclear degrees of freedom must be specifically tuned by the protein sequence to maintain the electronically excited state long enough to allow photon emission.

Enzyme catalysis is another key area in which there is a high degree of interplay between electronic and nuclear dynamics. There are numerous equilibrium and non-equilibrium aspects, including substrate binding, transition state stabilization, preservation of intermediates, and product release. In that regard, the mFAP/DFHBI complex is an ideal model system for studying a subset of those steps, including how proteins stabilize ligands in a particular conformation, and maintain a conformation for a long enough time for the desired electronic transition to take place.

Conventional quantum mechanics/molecular mechanics (QM/MM) have been enormously successful in modeling many aspects of protein-ligand interactions. Furthermore, time resolved femtosecond x-ray crystallography has provided insight into the average behavior of chromophore photoswitching^25^. However, as highlighted in this study, different regions of a rough protein-ligand energy landscape can lead to dramatically different behaviors of the excited states. This necessitates the need for extensive sampling of the ground state to approximate the Boltzmann ensemble that ultimately determines protein function.

## Methods

### Quantum Mechanics Simulations

To develop the potentials used to parameterize the molecular mechanics simulations, we carried out quantum mechanical simulations using the Gaussian 16 quantum chemistry package^26^. The equilibrium gas-phase geometry of the ground state was determined by density functional theory (DFT) using the B3LYP functional and the 6-31G basis. The gas-phase geometry of the excited state was found using time dependent density functional theory (TD-DFT) using the B3LYP functional and the 6-31G basis. We then calculated potential energy surfaces for both geometries by performing a rigid scan (keeping all other internal coordinates fixed) of the phenyl and imidazole dihedral angles at 10 degree intervals.

Our quantum mechanical calculations are limited in some important ways. The 6-31G basis is quite modest in size, especially given the anionic character of DHFBI that would typically require the use of diffuse basis functions. Moreover, the modest size of the basis can also limit the quality of the excited states obtained via TD-DFT. We expect, however, the excited state of interest to be valence in nature and therefore the use of a modest basis may prove sufficient to obtain results with semi-quantitive accuracy.

To check the magnitude of error that we can expect from using this basis, we ran the same DFT and TD-DFT calculations on geometries used in the previous rigid scan with a 6-311G+(2d,p) basis. With this basis, we found that the barrier to rotation in the excited state for the imidazole and phenyl dihedrals were lowered by 2.7 kJ/mol and 2.4 kJ/mol, respectively. (Figure S6) We thought those errors, in comparison to the estimated barrier heights (19 kJ/mol and 11 kJ/mol), should not preclude getting results with semi-quantitative accuracy.

Another important limitation of our calculations, as discussed above, is the inability of DFT and TD-DFT to accurately describe the avoided crossing region. This, however, is not the focus of this work where only the regions of the potential energy surface in the vicinity of the planar structure are of interest.

### DFHBI Ligand Parameterization

Generalized Amber force field (GAFF)^22^ molecular mechanics (MM) parameters for the phenolate form of DFHBI were created with the antechamber, parmchk, and tleap programs in Amber 16^27^. Those parameters were then converted to Gromacs format with ParmEd. To reparameterize the imidazole and phenyl dihedral angles using gas-phase QM energies, the geometry of the DFHBI was also optimized in vacuum with Gromacs^28^. Both dihedral angles were sampled one at a time starting from planar and iteratively moved by 10° increments, allowing all other degrees of freedom to relax via energy minimization. A given dihedral angle was enforced with two restraints, one on each ring atom that could be used to define the dihedral. The force constant on each restraint was 10,000 kJ mol^-1^ rad^-2^, with the only difference between the two restraints being a 180° phase shift in the target angle. In GAFF and other molecular mechanics force fields, the dihedral angle energy is defined:

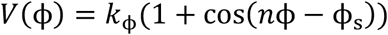

The dihedral angle is *φ* and three parameters are the multiplicity (*n*), force constant (*k_φ_*), and start angle (*φ*_s_). With *φ*_s_ = 180°, as is the case for all DFHBI dihedral angles, this function has a minimum at 0 and has *n* maxima starting at 180°/*n*. In the GAFF parameters for DFHBI, only single function was used for each dihedral. To model the quantum mechanical energies, we used a Fourier series of either four cosines for the ground state (n = 2, 4, 6, 8) or six cosines for the excited state (n = 2, 4, 6, 8, 10, 12). The free parameters in the optimization were the different *k_φ_*, one associated with each value of *n*. Prior to force constant optimization, the MM energies for each ligand conformation (excluding the four dihedral and dihedral restraint energies) were subtracted from the QM energies. The residual QM energies, which were assumed to represent only the contribution of the dihedral angle, were renormalized to start with 0 kJ mol^-1^ at 0°. The *k_φ_* were then optimized via linear least squares, using the observed dihedral angle values for each MM optimized ligand conformation. To avoid introducing artifacts in the fit potential energy surface from regions far from the metastable state, the fitting was only done on angles up to and including 80°. The iterative MM geometry optimization was then repeated. The *k_φ_* values converged after two rounds of parameter optimization, with the third round giving force constants to within 4-6 decimal places of the second round. The resulting parameters, which were used for ground and excited state simulations, are given in Table S1. The QM and reparametrized MM energies, along with the basis cosines, are shown in Figure S1.

The MM energies were parameterized assuming that the dihedral angles were energetically independent and could be treated in an additive manner. To determine whether that was the case, we repeated the QM geometry optimization, simultaneously restraining both dihedral angles on a 2D grid from 0-180° in 10° increments. (Figure S2) The ground state energies were very well approximated by an additive model, with significant deviations only occurring in inaccessible regions (> 200 kJ mol^-1^). The excited state energies were also well approximated by the additive model within the metastable basin (i.e. within a region about 30-40° of planar. However, the additive model did show a false minimum when both dihedrals were equal to 90°, which is close to the global maximum in the non-additive model. However, that region was not relevant for determining rotation kinetics. In practice, we found that the ~30 kJ mol^-1^ barrier in the additive model when both angles were around 40° (Figure S2B) was sufficient to direct the vast majority of simulation trajectories over a transition state barrier involving rotation of a single dihedral angle.

### Fluorescence Normalization and Selection of mFAP Pairs

Data from four different fluorescence experiments on the mFAP variants^5^ were normalized then averaged to determine a single fluorescence value. The four experiments consisted of combinations of two different final DFHBI concentrations (100 nM and 10 μM) and two different amounts of small-scale purified proteins (25 and 50 μL) in a total volume of 200 μL. For those experiments, the excitation and emission wavelengths were 488 and 510 nm, respectively. Raw fluorescence values were normalized by dividing each by the root mean square (RMS) fluorescence for the particular experiment they came from. The normalized fluorescence value used in this manuscript was the mean of the four values for a given variant, with the standard error calculated using the standard deviation divided by the square root of 2, to account for what was likely fewer independent experiments.

The sequences of those variants were aligned with Clustal Omega^29^ and only variants that aligned onto mFAP-A without gaps or insertions were retained. In addition, any variant that showed a low level of expression (protein gel density less than the 15 kDa ladder band) was eliminated. From the remaining variants, pairs were identified that differed by only a single amino acid substitution and showed at least a twofold difference in normalized fluorescence. This left 13 variants, with nine total pairs differing by a single amino acid. They are labeled mFAP-A through mFAP-M, in order from highest to lowest normalized fluorescence. The variant names, normalized fluorescence values, Kd values and gel densities are listed in Table S2.

### Preparation of mFAP Models

Unlike the initial mFAP designs^1^, structures for the variants analyzed in this study have not yet been determined. Furthermore, the designs include a five-residue extension of loop 7. Therefore, all simulations were started from models derived from the backbone of a designed model of mFAP-A. To sample a broad array of possible ligand conformations, we used 200 different DFHBI poses derived from docking to a designed model of mFAP1. Those ligand conformations were transferred onto to the mFAP-A structure by superimposing the Cα atoms of the mFAP1 binding site residues (14, 15, 16, 18, 22, 23, 24, 25, 26, 27, 28, 42, 43, 44, 45, 46, 50, 52, 72, 75, 78, 98, and 104) onto the equivalent residues of mFAP-A via sequence alignment. The ligand poses included all four major permutations of the ligand structure (i.e. from combinations of 180° rotations about the major and minor ligand axes).

To determine whether the sequence variations might affect the bound pose of DFHBI, we repacked the sequence of each variant around all 200 models. We used a three-stage protocol to first repack the mFAP sequence, minimize the side chains while allowing rigid body movement of DFHBI, then do an additional repack. This was implemented using RosettaScripts^30^, with the full XML protocol given in supporting methods. Of the four possible ligand conformations, we found that the lowest energy poses were the same as the original designed model and the mFAP1 crystal structure (6CZI). Therefore, for each variant we selected models derived from the first 20 ligand poses in that canonical conformation to start molecular dynamics simulations.

### Ground State Molecular Dynamics Simulations

Ground state molecular dynamics simulations were run with Gromacs 2019 using the Amber 99SB*-ILDN force field^31^, SPC/E water, and a NaCl concentration of 150 mM with Joung ions^32^. A dodecahedral box was sized with a 10 Å minimum distance between atoms in the protein/DFHBI and the edge of the box. The simulations were run at 300 K using a velocity rescaling thermostat^33^ and 1 bar using a Berendsen barostat^34^. A timestep of 2 fs was used and all bond lengths were constrained with the LINCS algorithm^35^. After minimization, the system was equilibrated using four successive 50 ps simulations with protein heavy atoms restrained to the starting conformation with the respective force constants: NVT (1000 kJ mol^-1^ nm^-2^), NPT (1000 kJ mol^-1^ nm^-2^), NPT (200 kJ mol^-1^ nm^-2^), and NPT (40 kJ mol^-1^ nm^-2^). Production ground state simulations were run for 100 ns and snapshots of all atomic positions and velocities were recorded every 50 ps.

### Excited State Molecular Dynamics Simulations

Excited state simulations were started from ground state simulation snapshots, preserving all atomic coordinates and velocities. The absorption of a photon was simulated by an instantaneous change of the dihedral angle potential for the imidazole and phenyl dihedrals using the MM potential shown in Figure S1, with parameters given in Table S1. All other simulation parameters were kept the same. Coordinates for DFHBI were saved every 50 fs. The DFHBI dihedral angles were analyzed on-the-fly every 10 seconds of wall-time using gmx gangle. When either the imidazole or phenyl dihedral angle exceeded 60°, the simulation was terminated and the time of rotation recorded.

Simulated quantum yields for the excited state DFHBI/mFAP simulations were also calculated using the observed Spinach-DFHBI fluorescence lifetime (τ_fluor_ = 4 ns) normalized by the quantum yield (QY = 0.72): τ_fluor_ = τ_fluor,obs_/QY = 5.56 ns. Assuming that fluorescence emission follows single exponential kinetics, the theoretical probability of emitting a photon for an excited state simulation with a given time in the planar state (*t*_planar_) is 1 − *e*^−*t*_planar_/*τ*_fluor_^. This formula was used to calculate the theoretical quantum yields in Figure S10, with the standard error determined from the 20 per-set average quantum yields.

### Quantum Mechanics Simulations with Relaxed Geometries

We calculated a relaxed potential energy surface, where other degrees of freedom are fully optimized aside from those being constrained, using the Gaussian 16 quantum chemistry package^26^. The constrained optimization was carried out using standard options in Gaussian which enforce the constraints via projection of the relevant degrees of freedom. In these relaxed scan calculations we used the 6-311G(2d,p) basis to address some of the limitations of the original basis set used. In the case of the excited state surface, we produced both a vertical excitation surface (using the relaxed ground state geometries) and an adiabatic excitation surface (with relaxed geometries in the excited state). The heights of the excited state energy barriers for 6-311G+(2d,p) rigid rotation and 6-311G(2d,p) relaxed rotation were closer to one another than the 6-31G rigid rotation, suggesting that the change of basis was primarily responsible for the lowering of the energy barrier, not the structural relaxation. (Figure S6) However, relaxation is likely responsible for shifting the imidazole energy barrier from 40° to 50°.

### ^26^DFHBI Ligand Reparameterization

After relaxing the ground and excited state DFHBI geometries with higher-level QM calculations, we then used a modified version of the parameterization protocol described above. We sampled the full two-dimensional grid of imidazole and phenyl dihedral angles as shown in Figure S8 (imidazole: 0-90° and phenyl 0-180°). During these scans, we found that the methine hydrogen on the linker could be in one of two positions above or below the plane of DFHBI, so both positions were tried during MM optimization and whichever gave the lowest MM energy was used. In addition, we enabled LINCS constraints on all bonds during MM energy minimization to mimic their use during the MD simulation. Because trans-DFHBI conformations (i.e. imidazole angles more than 90° from planar) were not used for MM parameterization, nor visited prior to jumping over the excited state energy barrier, those energies were not calculated.

Once the 2D potential energy surfaces were calculated, we then used 1D potential energy surfaces (i.e. with the other dihedral angle fixed at 0°) to parameterize the Fourier series of cosines in the ground and excited states for both the imidazole and phenyl dihedral angle potentials. We then calculated the difference between the 2D MM potential energy surface and the 2D QM potential energy surface, using the difference to create a CMAP correction^36^ to model the non-additivity of both dihedrals rotating simultaneously. The resulting 2D MM potentials very closely matched those shown in Figure S8.

## Acknowledgements

The authors thank Jason Klima for many helpful discussions and advanced access to the new mFAP designs, functional characterization, and docked DFHBI conformations. This work was supported by a National Science Foundation XSEDE Research Allocation (MCB190110).

