## Supplemental Information for "Prediction of fluorophore brightness in designed mini fluorescence activating proteins"

### Supporting Methods

#### Preparation of mFAP Models

The following RosettaScript was used for repacking the mFAP sequences around the DFHBI ligand conformations. The input files were a PDB file, a resfile with the mFAP sequence, and a file called HBI.fa.params with parameters for the HBI ligand. It was run with a command line similar to the following:

```
rosetta_scripts -s *.pdb -parser:protocol energyminimization.xml -
packing:resfile mFAPX.resfile -extra_res_fa HBI.fa.params
```

```
<ROSETTASCRIPTS>
	<SCOREFXNS>
		<ScoreFunction name="r15" weights="ref2015" />
	</SCOREFXNS>
	<RESIDUE_SELECTORS>
	</RESIDUE_SELECTORS>
	<TASKOPERATIONS>
		<ExtraRotamersGeneric name="extrachi" ex1="1" ex2="1"/>
		<RestrictToRepacking name="repackonly"/>
		<ReadResfile name="mFAP"/>
	</TASKOPERATIONS>
	<MOVE_MAP_FACTORIES>
	</MOVE_MAP_FACTORIES>
	<SIMPLE_METRICS>
	</SIMPLE_METRICS>
	<FILTERS>
	</FILTERS>
	<MOVERS>
		<PackRotamersMover name="repack1" nloop="5" scorefxn="r15"
			task_operations="mFAP,extrachi"/>
		<MinMover name="minimize1" scorefxn="r15" chi="true" bb="false"
			jump="ALL" tolerance="0.001">
		</MinMover>
		<PackRotamersMover name="repack2" nloop="5" scorefxn="r15"
			task_operations="extrachi,repackonly"/>
	</MOVERS>
	<PROTOCOLS>
		<Add mover="repack1"/>
		<Add mover="minimize1"/>
		<Add mover="repack2"/>
	</PROTOCOLS>
	<OUTPUT scorefxn="r15"/>
</ROSETTASCRIPTS>
```

#### Supporting Figures

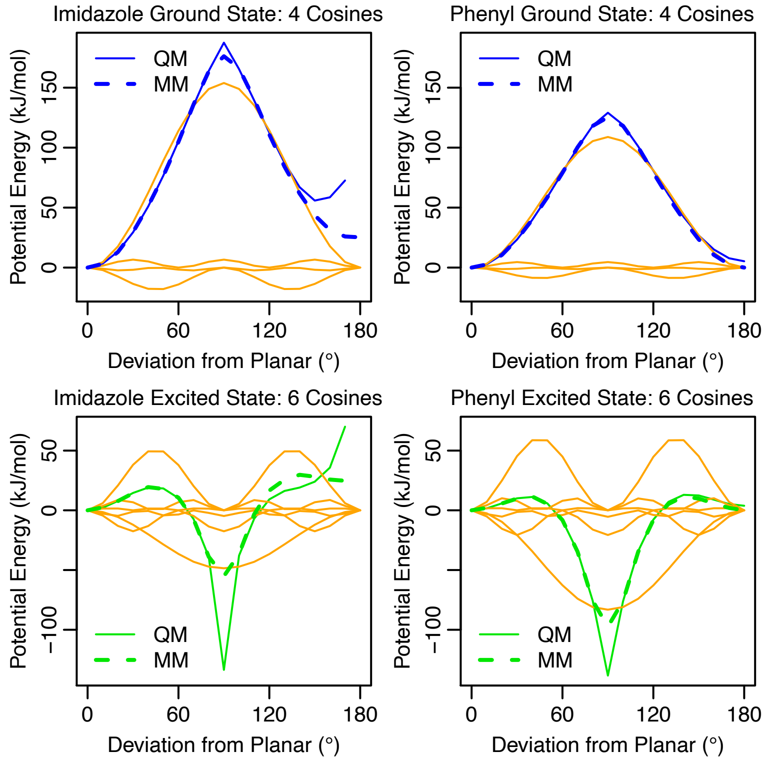


**Figure S1. Quantum mechanical energies modeled with a series of cosine functions.** Quantum mechanical energies (QM, solid lines) are match molecular mechanics energies (MM, dashed lines) modeled sum of basis cosine functions (orange lines). Only angles less than or equal to 80° were used to optimize the force constants. The force constants are given in Table S1.

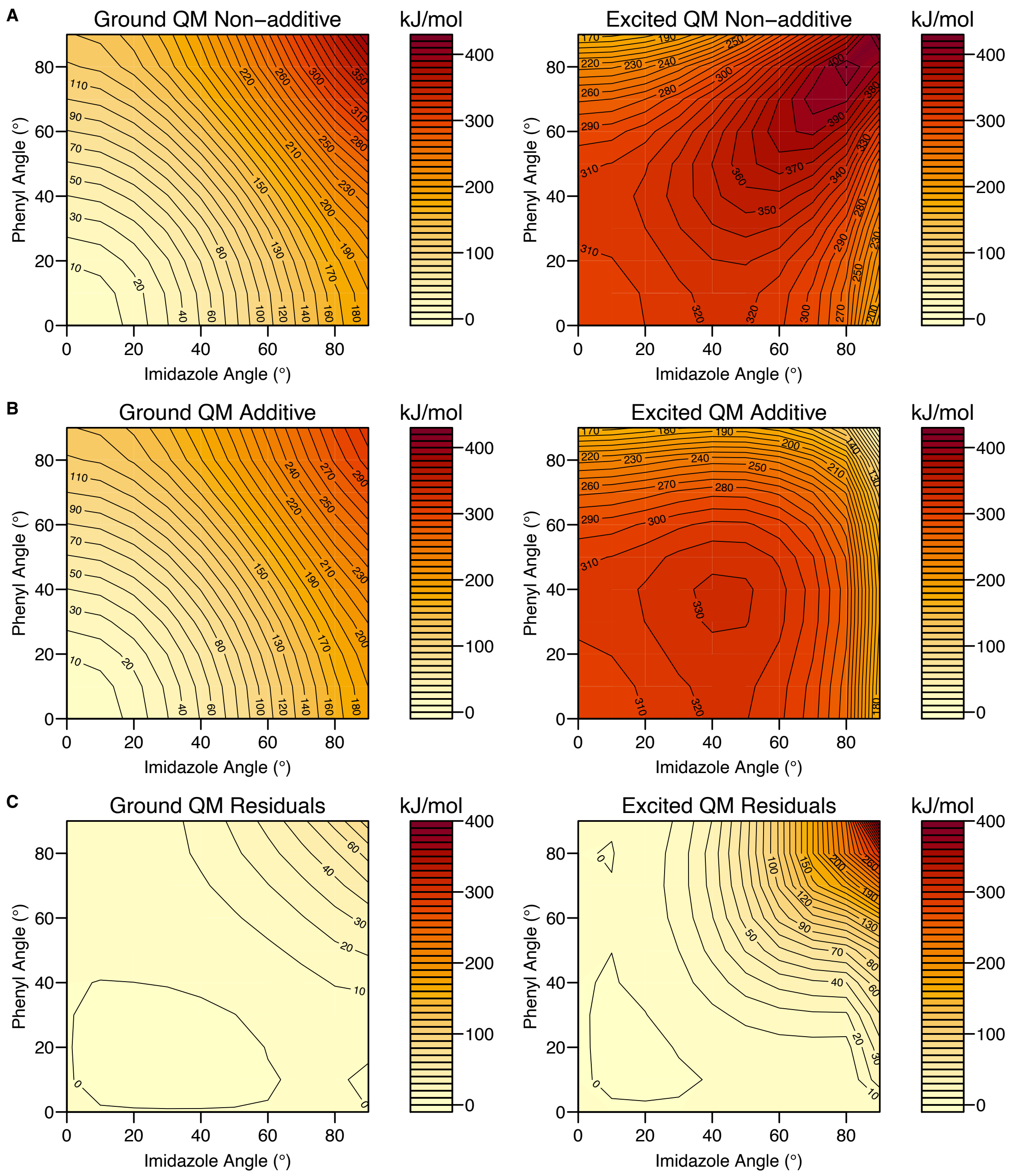


**Figure S2. Additive potential approximates non-additive potential in metastable region.** **A)** Quantum mechanical (QM) energies for the ground and excited states sampled on a 2D grid with both angles restrained simultaneously. **B)** An additive model assumes $E(\varphi_{\text{imidazole}}, \varphi_{\text{phenyl}}) = E(\varphi_{\text{imidazole}}, 0) + E(0, \varphi_{\text{phenyl}})$, similar to the molecular mechanics parameterization. **C)** The two models are very close in the metastable region (within about 30-40° of planar).

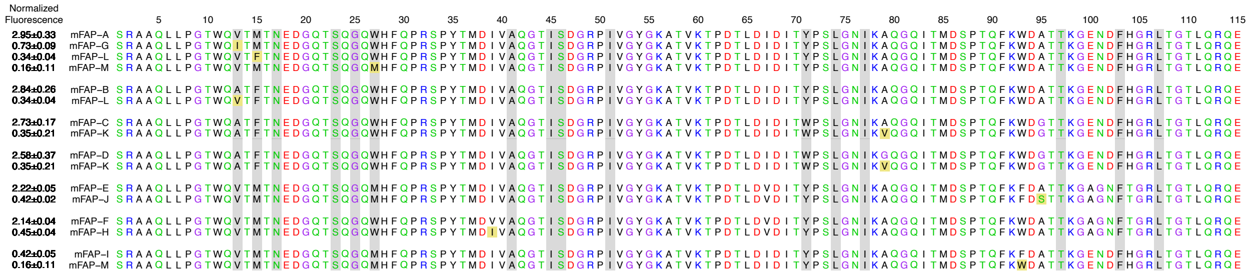


**Figure S3. mFAP variants used in this study.**

In each group of sequences, there is a single amino acid difference (yellow) between the top sequence and the sequences below. Residues that interact with the DFHBI ligand are highlighted in gray.

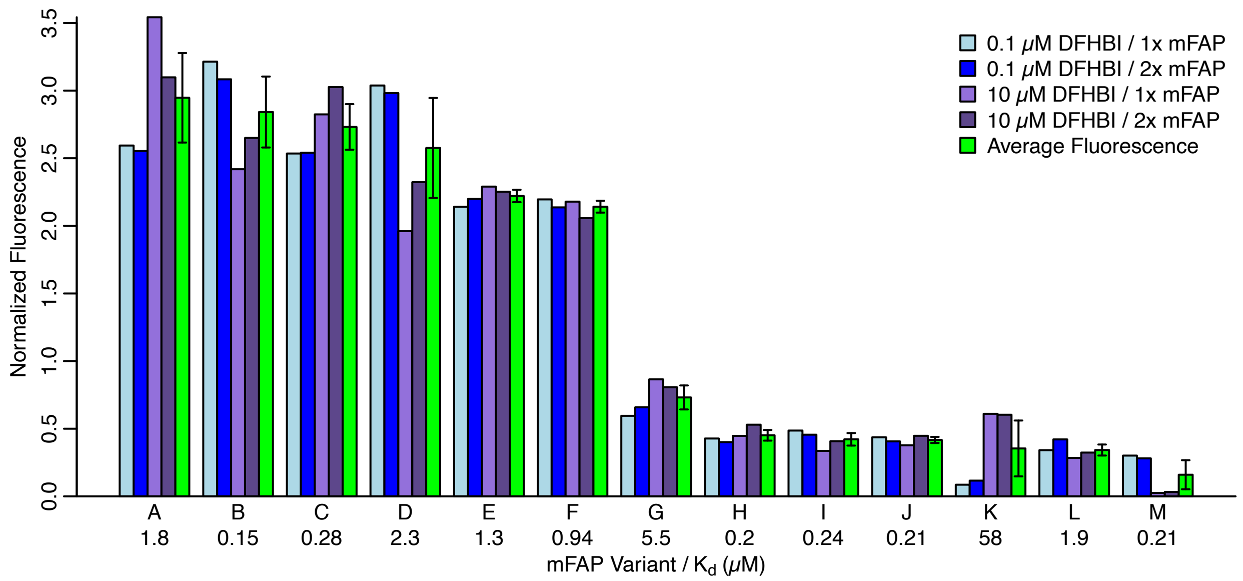


**Figure S4. Calculation of average normalized fluorescence.**

A total of four experiments were used to measure fluorescence from small-scale expression. They used combinations of either a 0.1 or 10 μM final concentration of DFHBI, and a 1x (25 μL) or 2x (50 μL) volume of added mFAP. The fluorescence values for all 92 variants were normalized by dividing the root mean square (RMS) fluorescence from the particular experiment. The average normalized fluorescence, used in this manuscript, is shown in green. Several variants with weaker binding affinities ($K_d > 1$ μM) show reduced observed fluorescence at the lower DFHBI concentration, which may be a result of incomplete mFAP/DFHBI binding. Those variants include mFAP-A, mFAP-G, and especially mFAP-K. mFAP-M shows unexpectedly low fluorescence at the higher DFHBI concentration, possibly causing a downward bias in its normalized fluorescence.

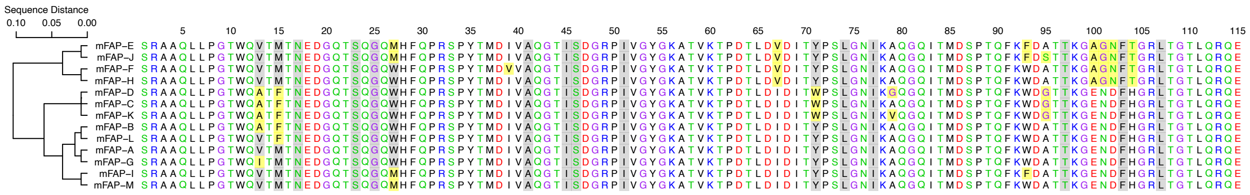


**Figure S5. mFAP variants clustered by sequence.**

mFAP sequences were clustered by sequence using Clustal Omega. Amino acid positions differing from mFAP-A are highlighted in yellow. Residues that interact with the DFHBI ligand are highlighted in gray.

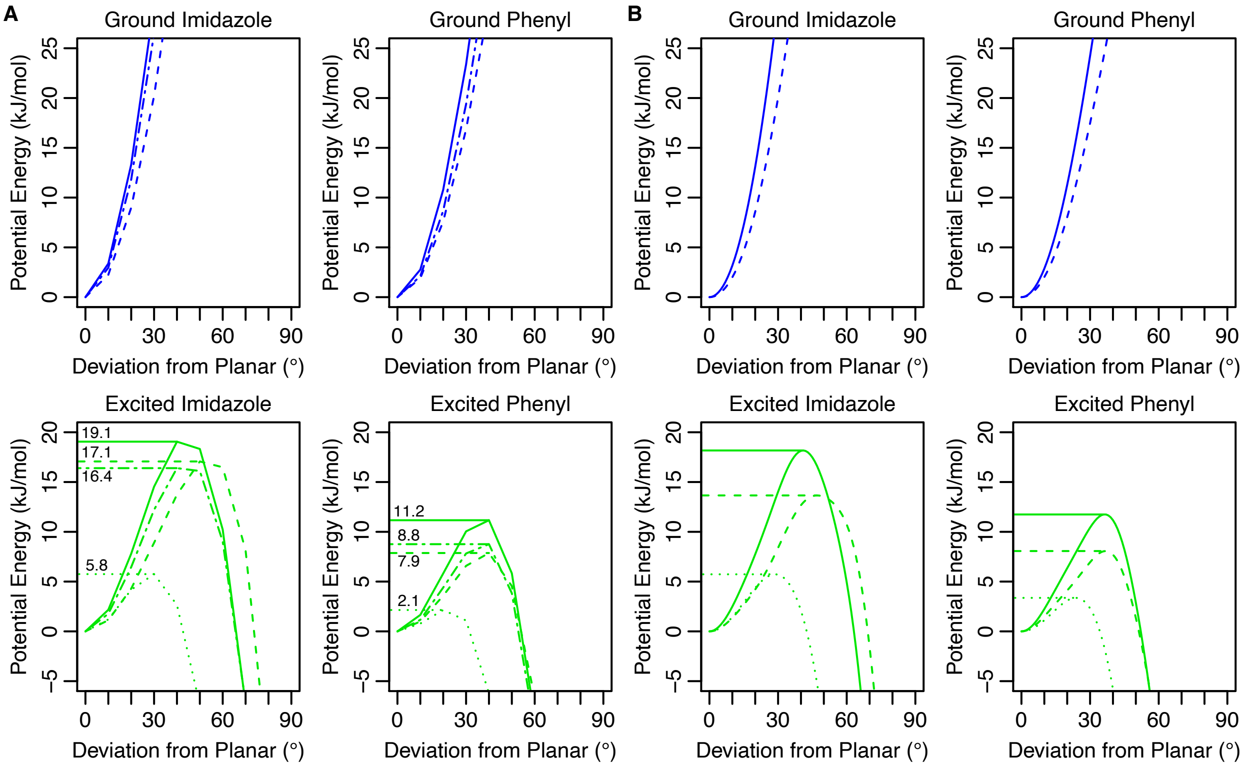


**Figure S6. 1D QM and MM potential energies with and without coordinate relaxation.** **A)** QM potential energy surfaces calculated without coordinate relaxation (i.e. using rigid dihedral rotations from geometries relaxed in the ground or excited states) using 6-31G basis are shown with solid lines. They are the same as values shown in Figure 1, Figure S1, and Figure S2. Potential energy surfaces calculated without coordinate relaxation using 6-311G+(2d,p) basis are shown with dotted-dashed lines. Potential energy surfaces calculated with coordinate relaxation in the ground state using 6-311G(2d,p) basis are shown with dashed lines and are the same as shown in Figure S8A/B. Potential energy surfaces calculated with coordinate relaxation in the excited state using 6-311G(2d,p) basis are shown with dotted lines and are the same as shown in Figure S8C. To aid in visualizing differences between the energy barriers, horizontal line segments are drawn from the peak of the excited state energy profile to the y-axis. **B)** The MM dihedral angle potentials parameterized to fit the respective QM energies. Differences between the energies in A and B come from non-dihedral terms in the MM energy for DFHBI.

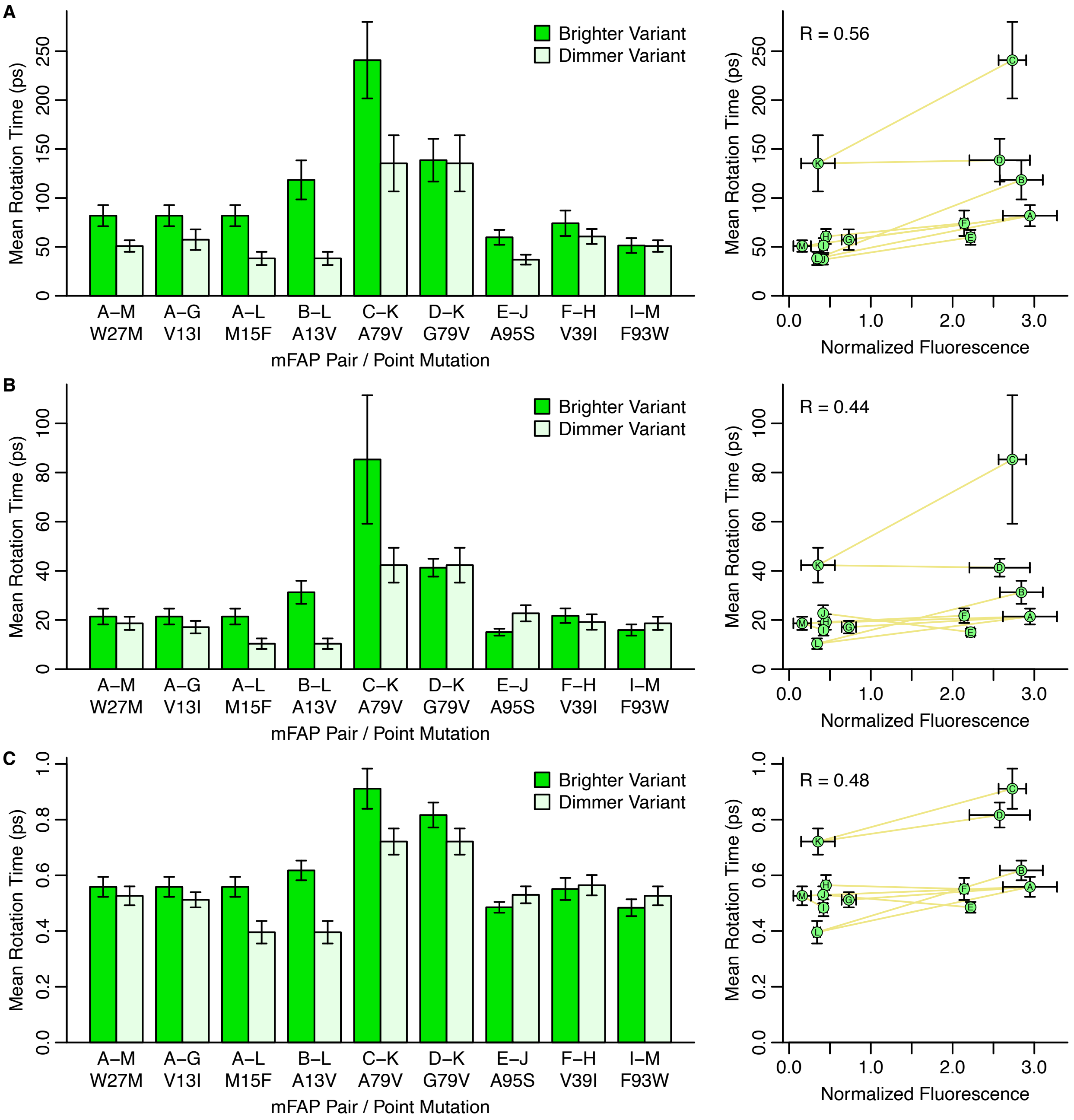


**Figure S7. Mean rotation times using different ground and excited state potentials.**

**A)** Rotation times using MM energies parameterized from rigid rotation of optimized planar geometry (Figure S6, solid lines). Data is the same as shown in Figure 3C and Figure 4A **B)** Rotation times using MM energies parameterized from geometries optimized in the ground state (Figure S6, dashed lines). **C)** Rotation times using MM energies parameterized from geometries optimized in the ground and excited states (Figure S6, blue dashed lines then green dotted lines).

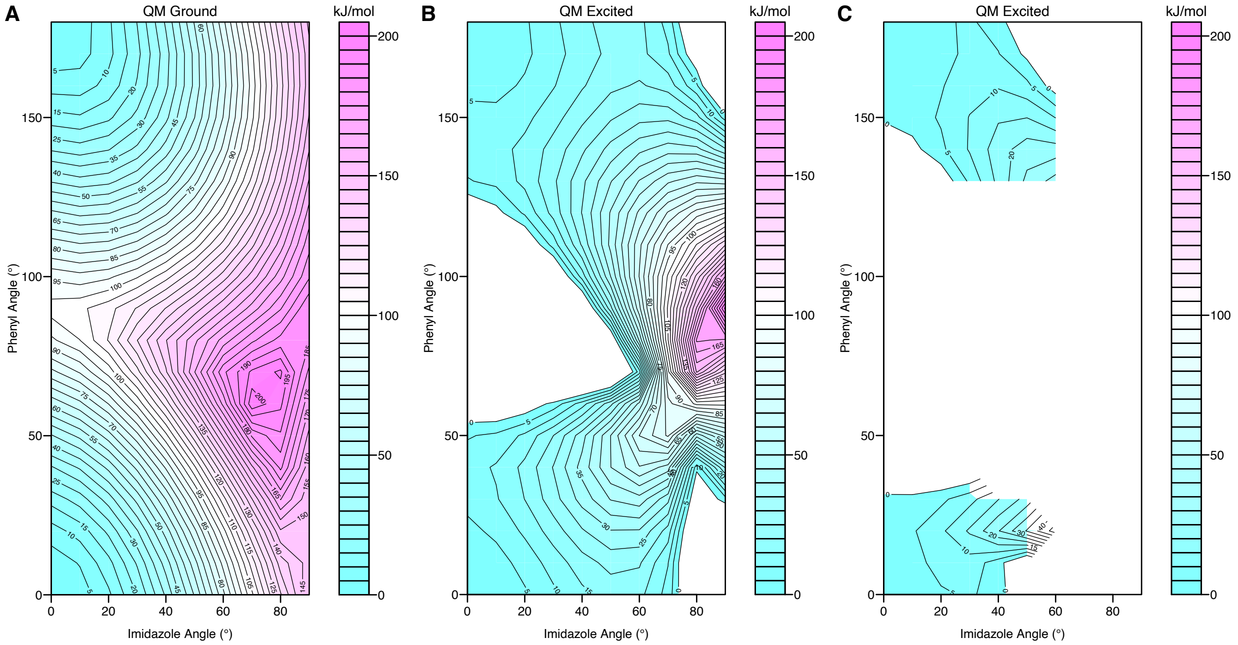


**Figure S8. 2D QM potential energy surfaces using coordinate relaxation.**

Potential energy surfaces are shown for the imidazole and phenyl dihedrals. The full potential energy surfaces are created by applying symmetry operations. All potential energy surfaces are set to 0 kJ/mol at angles of 0° and 0°. **A)** Ground state energies allowing relaxation of all other degrees of freedom. **B)** First excited state energies using the same geometry used in **A**. **C)** First excited state energies allowing relaxation of all other degrees of freedom. Energies were only calculated for imidazole angles 0-60° and phenyl angles 0-40° and 120-180°.

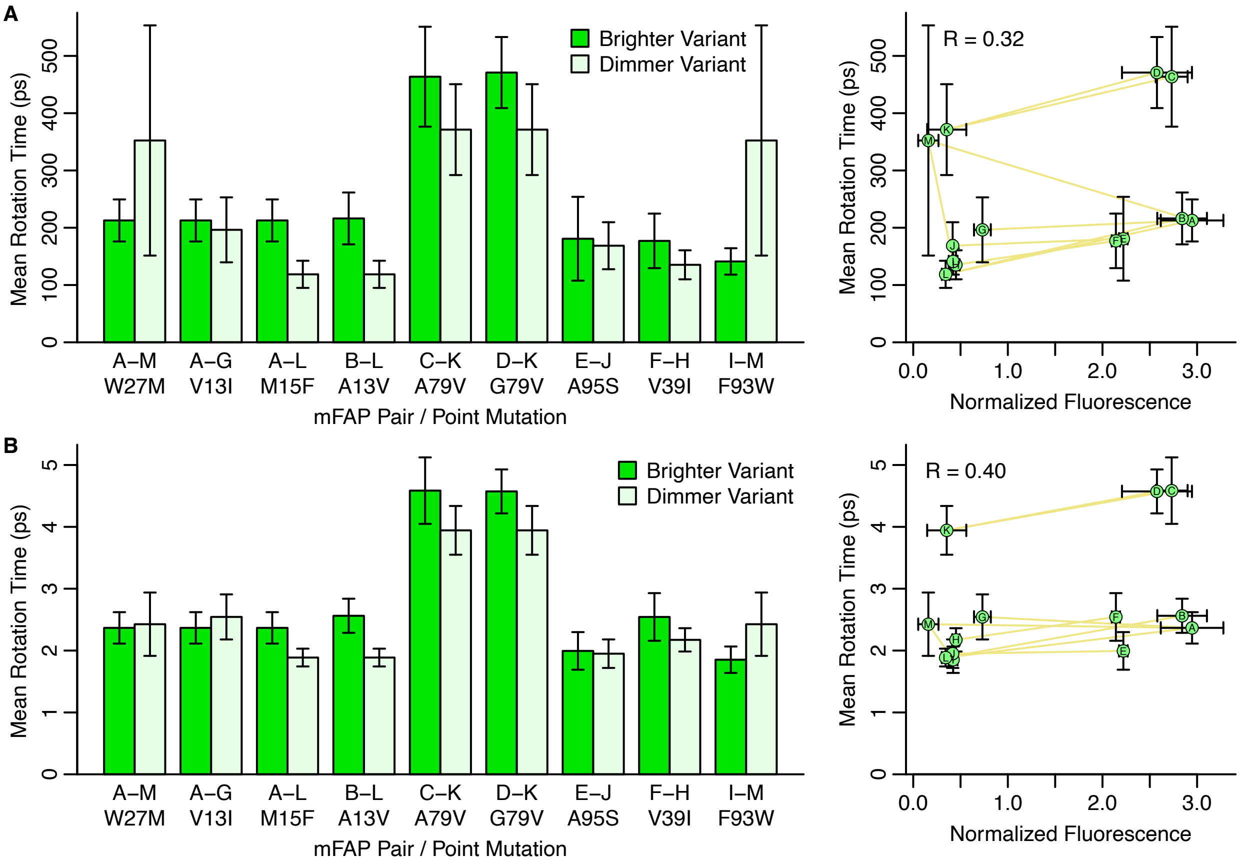


**Figure S9. Mean rotation times from 2D quantum mechanical potential energy surfaces.** The ground state was simulated using a molecular mechanics 2D potential energy surface fit to the quantum mechanical energies shown in Figure S8A. **A)** Mean chromophore rotation times from excited state simulations using a 2D potential energy surface parameterized using vertical excitations as shown in Figure S8B. **B)** Mean chromophore rotation times from excited state simulations using a 2D potential energy surface parameterized using relaxed excited state simulations as shown in Figure S8C.

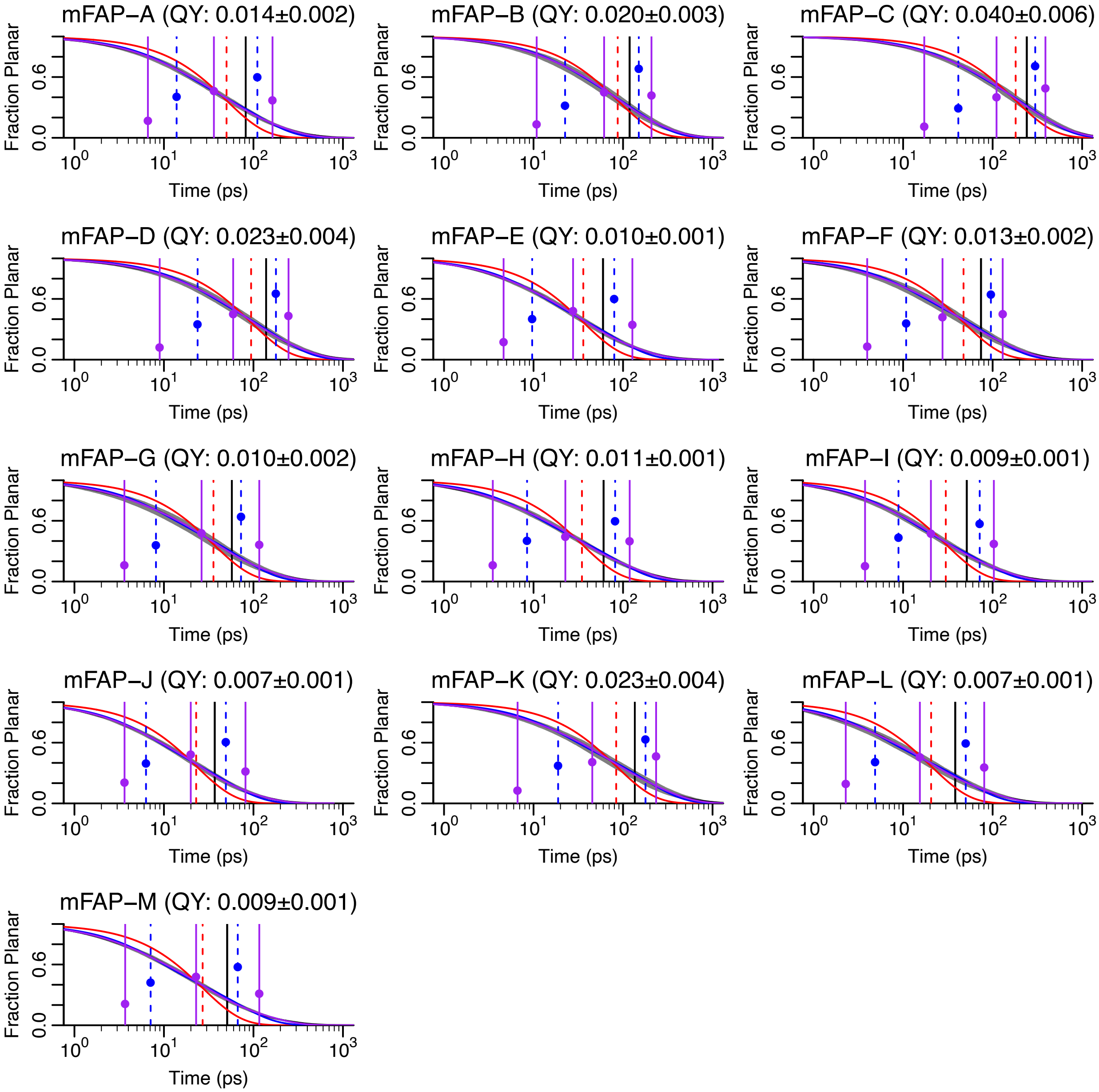


**Figure S10. Multiexponential escape kinetics in trajectories of mFAP/DFHBI.**

For each mFAP variant, the average planar decay curve (black) and standard error (shaded gray) is shown along with the average lifetime (vertical black line). Single (red), double (blue), and triexponential (purple) fits are to the equation $f(x) = \sum c_i e^{-x/\tau_i}$ with the constraint $\sum c_i = 1$. $\tau_i$ values are shown as vertical dashed lines and $c_i$ values are shown as filled points. Quantum yields are calculated assuming $\tau_{fluor} = \tau_{fluor,obs}/QY = 5.56$ ns.

#### Supporting Tables

| | Imidazole Dihedral | | Phenyl Dihedral | |
|---|---|---|---|---|
| | $n$ | $k_\varphi$ (kJ mol$^{-1}$) | $n$ | $k_\varphi$ (kJ mol$^{-1}$) |
| Ground State | 2 | 38.4753 | 2 | 27.2273 |
| | 4 | -4.5921 | 4 | -2.2127 |
| | 6 | 1.6565 | 6 | 1.1532 |
| | 8 | -0.5933 | 8 | -0.3248 |
| Excited State | 2 | -12.1275 | 2 | -20.7976 |
| | 4 | 12.7229 | 4 | 15.1414 |
| | 6 | -4.3846 | 6 | -5.1656 |
| | 8 | 2.2351 | 8 | 2.6065 |
| | 10 | -1.1894 | 10 | -1.3605 |
| | 12 | 0.4948 | 12 | 0.5725 |

**Table S1. Ground and excited state energy parameters.**

The equation used to evaluate these parameters was $V(\phi) = k_{\phi}(1 + cos(n\phi - \phi_s))$, with φ being the dihedral angle. The $\varphi_s$ parameter was set to 180° for all dihedrals. Graphs of the individual dihedral potential energy functions resulting from these parameters are shown as dashed lines in Figure S1 and solid lines in Figure S6B.

| Variant | Klima ID[5] | Normalized Brightness | DFHBI $K_d$ (μM)[5] | Gel Density[5] |
|---|---|---|---|---|
| mFAP-A | mFAP2b | 2.95±0.33 | 1.800±0.250 | 5.2 |
| mFAP-B | mFAP2a | 2.84±0.26 | 0.150±0.011 | 4.2 |
| mFAP-C | mFAP4 | 2.73±0.17 | 0.284 | 4.8 |
| mFAP-D | mFAP3 | 2.58±0.37 | 2.262±1.220 | 3.1 |
| mFAP-E | mFAP2.4 | 2.22±0.05 | 1.314 | 8.1 |
| mFAP-F | mFAP2.2.9 | 2.14±0.04 | 0.945 | 10.4 |
| mFAP-G | mFAP2.5.5 | 0.73±0.09 | 5.526 | 5.8 |
| mFAP-H | mFAP2.2.13 | 0.45±0.04 | 0.200 | 10.3 |
| mFAP-I | mFAP_pH | 0.42±0.05 | 0.242±0.039 | 2.4 |
| mFAP-J | mFAP2.2.15 | 0.42±0.02 | 0.206 | 7.7 |
| mFAP-K | mFAP8 | 0.35±0.21 | 58.123 | 3.5 |
| mFAP-L | mFAP2a.0 | 0.34±0.04 | 1.898 | 3.4 |
| mFAP-M | mFAP2c.1 | 0.16±0.11 | 0.213 | 2.6 |

**Table S2. mFAP variants simulated.**

Gel density is relative to a 15 kDa protein ladder band from Precision Plus Protein Unstained Protein Standard (Bio-Rad)

Will Barr
Isabela Camacho-Horvitz
Devon B. Cooper
Nicole DelGaudio
Emma R. Hostetter
Meera Joshi
Jana C. O'Donnell
Ivy Poon
Elliot J. Williams

**Table S3. Students in 2018 Wesleyan Molecular Modeling and Design course.**
